# Autocorrelation analysis of a phenotypic screen reveals hidden drug activity

**DOI:** 10.1101/2023.03.14.532578

**Authors:** Richard A. Dubach, J. Matthew Dubach

## Abstract

Phenotype based screening is a powerful tool to evaluate cellular drug response. Using high content fluorescence imaging of simple fluorescent labels and complex image analysis, phenotypic analysis identifies subtle compound-induced cellular changes unique to compound mechanisms of action (MoA). Recently, a screen of 1,008 compounds in three cell lines was reported where phenotype analysis detected changes in cellular phenotypes and accurately identified compound MoA for roughly half the compounds. However, we were surprised that DNA alkylating agents and other compounds known to induce or impact the DNA damage response produced no activity in cells with fluorescently labeled TP53BP1 - a canonical DNA damage marker. We hypothesized that phenotype analysis is not sensitive enough to detect small changes in 53BP1 distribution and analyzed the screen images with autocorrelation image analysis. We found that autocorrelation analysis, which quantifies the clustering of fluorescently-labelled protein within the nucleus, of 53BP1 images from this screen identified higher compound activity for compounds and MoAs known to impact the DNA damage response. These results demonstrate the capacity of autocorrelation to detect otherwise undetectable compound activity and suggest that autocorrelation analysis of specific proteins could serve as a powerful screening tool for drug discovery.

## Introduction

Phenotypic screening is a potent tool to identify compounds that alter cellular function or properties (1). Historically, phenotypic screening has played a significant role in identifying many of the drugs that are currently in the clinic (2). More recent applications of phenotypic screening, or image based profiling (3), use high throughput fluorescence imaging of cells with fluorescent labels that capture the shape and structure of the cell and organelles (4). Morphological profiling of large datasets (5) enables assignment of compound mechanism of action by comparing morphological properties to known compounds or characterization of the impact of genetic perturbations (6). Because these screens don’t really on *a priori* knowledge of key targets, they can provide profound insight in the drug discovery pathway (7).

Recently a screen of 1,008 compounds generated mechanism of action (MoA) identification of compounds with unclear mechanisms through comparison to known MoAs (8). The screen included three different cells types that expressed combinations of fluorescently labeled proteins: ACTB-RAB5A, CANX-COX4I1, GM130-SQSTM1, TUBA1B-RELA, and TP53BP1-CLTA, generating 15 different cell lines. Phenotypic analysis extracted features from segmented cells to classify compound MoA based on unique MoA descriptors. The screen accurately ranked roughly half the testable MoAs of the reference compounds. Yet, a substantial number of MoAs did not produce identifiable activity. For all three cell lines, TP53BP1-CLTA labeled cells did not produce increased sensitivity to compounds with DNA damage, DNA damage response, or cell cycle MoAs. A surprising result considering 53BP1 is a canonical DNA damage response marker (9).

53BP1 is a component of the DNA double strand break response pathway and recruited to sites of DNA damage into foci that form around damaged DNA (10). There are myriad components that impact 53BP1 activity, including ATM activity (11), cell cycle (12) and epigenetic modifications (13). Therefore, compounds that induce DNA damage, alter the cell cycle, impact DNA signaling or alter the DNA damage response are expected to affect the nuclear location of 53BP1. In theory, any altered localization of 53BP1 would be identified by a phenotypic screen to reveal compound activity in cells with labeled 53BP1. However, this was not the case in the original analysis of the imaging data. Therefore, here we hypothesized that image autocorrelation analysis of 53BP1 images would reveal altered 53BP1 localization that was not detectable using traditional phenotypic screening. Image autocorrelation enables quantification of the spatial heterogeneity of fluorophores that is not possible with traditional analysis due to background noise present in all fluorescent imaging (14). Thus, autocorrelation provides a potentially more sensitive measurement of compound induced changes in 53BP1 localization.

## Results

We accessed the original phenotypic screen (8) images through the IDR API on the Open Microscopy server. The screen contained three different cell lines that had endogenous TP53BP1 labeled GFP as a marker protein. We first identified all the images from these three cell lines (>60,000 images in total), then segmented the nuclei of each cell using the BFP channel image. Segmented nuclear regions were then transferred to the TP53BP1 image and image autocorrelation was performed on each nucleus. The degree of aggregation (DA, a measurement of fluorophore clustering) for each nucleus was then averaged to produce an overall image 53BP1 DA that corresponded to the compound and compound concentration. The dataset contained repeats of four different doses for each compound. The activity of each compound in each cell line was then determined by fitting a linear regression to the DA as a function of compound concentration. The slope and significance versus the null hypothesis (slope equal to zero) of DA vs. concentration was then determined. Active compounds were defined as those having a significant (p<0.05) slope of DA vs. concentration.

53BP1 DA is a measure of protein labeled fluorophore clustering within the nucleus. A positive regression of DA vs. concentration indicates that the compound induces 53BP1 recruitment to foci at sites of DNA repair and/or processing. Increases in 53BP1 recruitment can occur either through induction of DNA damage or altered repair pathways, such as shifting the response from homologous recombination to non-homologous end joining. Conversely, a negative regression indicates that the compound prevents 53BP1 recruitment to DNA damage or reduces the amount of DNA damage in the cell. However, it should be noted that, as a phenotypic measurement, there are other potential mechanisms of altered 53BP1 clustering that could drive observed compound activity. Yet, given the highly characterized role of 53BP1 in the DNA damage response (9), recruitment to DNA damage response foci is likely the most prominent driver of measured activity.

Volcano plots for each cell line were generated to visualize the results (**Fig. 1**). A few compounds with significant activity and targets known to be involved in DNA handling, DNA damage response or cellular cycle are identified in the results. Table S1 contains the complete results. Overall, the majority of compounds with the strongest activity have mechanisms of action that impact the DNA damage response. Mirin, which inhibits Mre11 competition with 53BP1 at stalled replication forks (15) produced a strong response in HepG2. Bromodomain inhibitors impact 53BP1 signaling (16) - PFI 1 showed strong positive activity in both HepG2 and A549 cell lines, while PF CBP1, another bromodomain inhibitor, has strong activity in WPMY-1 cells. Other compounds that impact the DNA damage response were also strong inducers of 53BP1 recruitment. These include, in WPMY-1 cells: A66 (a p110α selective PI3K inhibitor), thiotepa (a DNA alkylating agent), and SAHA (a HDAC inhibitor).

**Figure 1.**
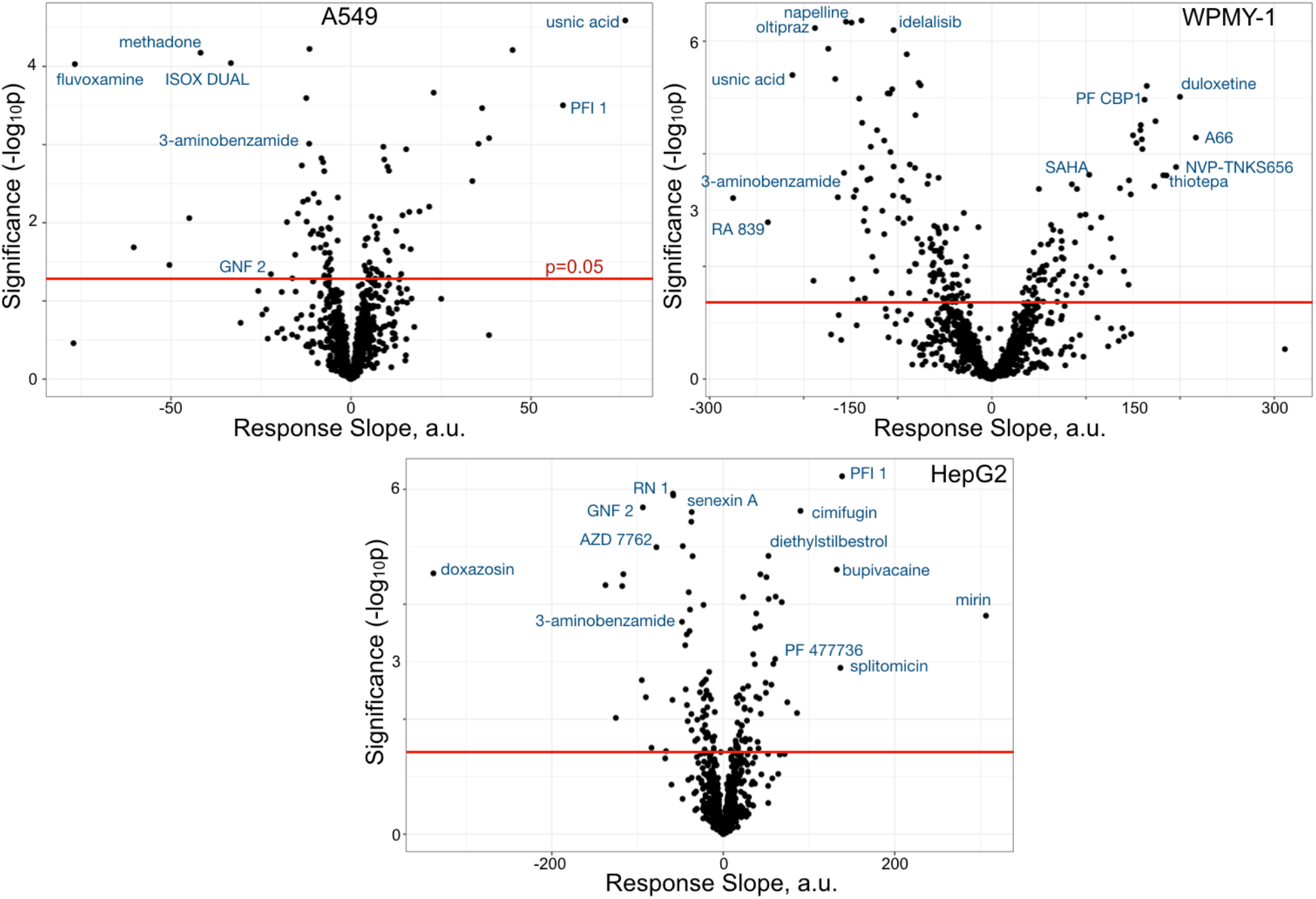
Compound activity on 53BP1 recruitment. Volcano plots for DA vs. concentration response slope linear regressions for each cell line. Above the red line (p=0.05) are compounds with significant activity. Identified compounds are known to impact the DNA damage response. Full results are in Table S1.

However, numerous compounds had significant activity reducing the recruitment of 53BP1. As an example, Nrf2 activators oltipraz and RA839 were two of the strongest 53BP1 recruitment-reducing compounds in WPMY-1 cells. Nrf2, a transcription factor, plays a role in the DNA damage response (17) and promotes homologous recombination repair (18), which reduces 53BP1 recruitment. Other compounds that impact DNA damage also have activity in our analysis. For example, the ABL1 inhibitor GNF 2 reduced 53BP1 recruitment in both HepG2 and A549 cells, likely through decreasing DNA damage (19). Curiously, usnic acid strongly induced 53BP1 recruitment in A549 cells but strongly prevented recruitment in WPMY-1 cells. However, the mechanism of action of usnic acid in the cellular DNA damage response remains unresolved (20), warranting further exploration of the differences between these cell lines.

We also determined a mechanism of action (MoA) activity score for each MoA containing results from at least 5 compounds through calculating the fraction of compounds with significant activity (**Fig. 2**). Compounds that prevent 53BP1 recruitment have a negative score while compounds that increased 53BP1 score positive. Thus, for MoAs with compounds that are both negative and positive, such as PARP inhibitors, the activity score is lower than the total number of active compounds. The MoAs that have the most activity in inducing 53BP1 DNA damage recruitment are ATM, GSK3 and MBT domain inhibitors, all compounds that canonically impact the DNA damage response. Conversely, some of the lowest scoring MoAs were ABL1 inhibitors, which reduce DNA damage (19), FAK inhibitors and LXR agonists, which both impact DNA repair without clear roles (21, 22), mechanistically suggesting that 53BP1 recruitment is reduced in cells treated with these compounds.

**Figure 2.**
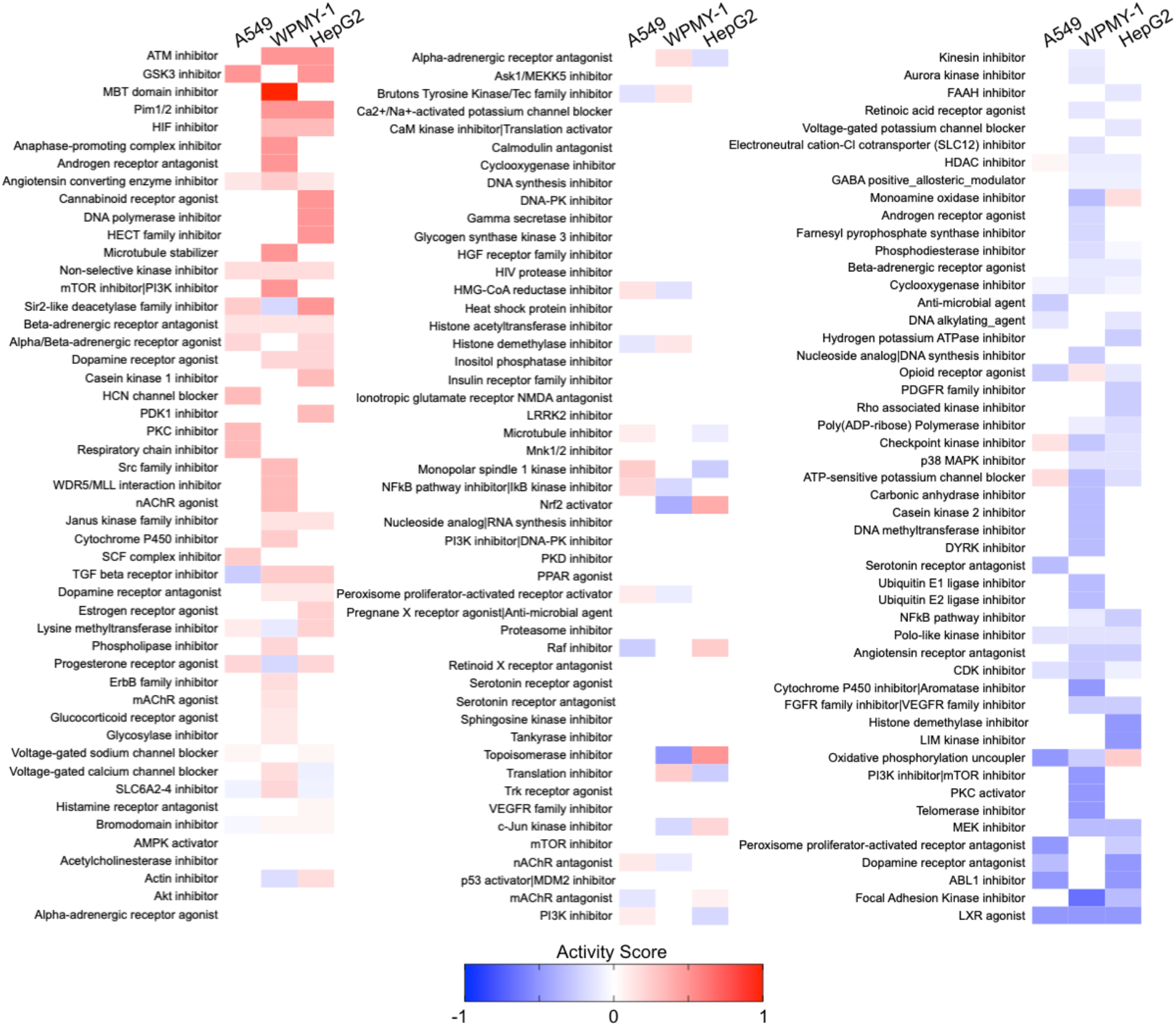
Mechanism of Action activity on 53BP1 recruitment to DNA damage. Activity scores for mechanisms of action with at least 5 compounds.

Some MoAs contained compounds that either reduced 53BP1 recruitment or increased recruitment. For example, the PARP inhibitor 3-aminobenzamide reduced 53BP1 recruitment in each of the 3 cell lines. However, the PARP inhibitors NVP-TNKS656, NU 1025, and 4-HQN increased 53BP1 recruitment in at least one cell line. Given the history of PARP inhibitors being misclassified (23) and the absence of more advanced clinical PARP inhibitors in the screen, these divergent results could stem from promiscuous or mis-identified compound MoAs.

In the original phenotypic analysis DNA alkylating agents had no measured activity while PARP inhibitors had very low activity - a surprising finding since both MoAs impact 53BP1 signaling (24, 25). However, our analysis found that DNA alkylating agents indeed have activity - 33% in WPMY-1, 25% in HepG2, and 11% in A549 cells - and PARP inhibitors have a higher activity than previously measured - 33% in WPMY-1, 15% in HepG2, and 17% in A549 cells. Overall, the increased activity observed in WPMY-1 cells could arise from higher 53BP1 signaling in these cells or aspects of the imaging and analysis. Unfortunately, images in the phenotypic screen have binned pixels. Binning serves to reduce the number of pixels over which the 53BP1 signal can be autocorrelated and increase the pixel size to limit spatial heterogeneity, which both impact the sensitivity of our analysis (26). In the images, HepG2 cells had the smallest nuclei, while A549 cell nuclei had a 15% bigger area and WPMY-1 cell nuclei were 40% larger.

Therefore, autocorrelation analysis of the WPMY-1 is expected to be more sensitive to changes in 53BP1 recruitment. Non-binned pixels would generate 4 times more pixels per nucleus with a quarter of the area, suggesting that non-binned images would generate greater analysis sensitivity.

## Discussion

In the original analysis of the imaging dataset no DNA alkylating agents compounds were found to be active. Furthermore, in A549 and WPMY-1 cell lines, the use of TP53BP1 as a marker actually reduced the ability to detect PARP inhibitor activity compared to other protein markers. This was similar for other MoAs that act on DNA or the DNA damage response, such as bromodomain inhibitors and HDAC inhibitors. Considering that 53BP1 is heavily involved in the DNA damage response pathway (27, 28) and many MoAs of the compounds screened act to interfere with the DNA damage response, alter the cell cycle or impact the amount of DNA damage in a cell, it was surprising that TP53BP1 was not a more sensitive marker in the phenotypic screen.

Yet, the recruitment of 53BP1 to sites of DNA damage is typically only resolved through pre-extraction and immunofluorescence (29). In this process, labile nuclear 53BP1 protein is extracted from the cell prior to fixation to reduce the background concentration and increase the resolution of chromatin-bound 53BP1 in DNA damage foci. The phenotypic screen analyzed here used live cells with fluorescently labeled, endogenous 53BP1, which prevents removal of protein not interacting with DNA and reduces the ability to resolve 53BP1 foci. This limitation likely prevents traditional phenotypic analysis from detecting non-resolvable spatial signaling of DNA damage response proteins. Furthermore, pre-extraction is a subjective process that potentially removes protein associated with chromatin and DNA and thus not a robust approach for phenotypic screening generally (30, 31).

Applying spatial image autocorrelation overcomes the limitation of high non-foci background fluorescence to quantify the degree of 53BP1 protein clustering within the nucleus. Here, we found autocorrelation analysis is able to detect compound activity of compounds that generated no activity when analyzed by traditional phenotypic analysis. These results confirm that many of the compounds associated with DNA damage, DNA damage response or cell cycle indeed have activity in the cell lines used in the screen. The results also demonstrate the power of spatial image autocorrelation of labeled specific proteins to quantify compound activity where traditional approaches have less sensitivity. Overall, these results suggest that more complex analysis of specific, yet broadly functional fluorescent labels can reveal compound activity that is not otherwise detectable.

## Methods

Images from the original study (8) were accessed through the Image Data Resource (32) API on the Open Microscopy environment. Nuclei were segmented using StarDist (33) on the BFP channel image. Segmented nuclei were then analyzed by image correlation spectroscopy autocorrelation (34) in the TP53BP1 channel and the average degree of aggregation was calculated. For each cell line, compound induced DA was plotted against the compound concentration and linear regression was performed with a linear model in R. MoA activity was determined for MoAs with at least 5 compounds with linear regression results in each of the cell lines. Active compounds were defined as having significant (p<0.05) non-zero linear regressions, either negative or positive. The activity score was determine by summing the direction of each active compound and dividing by the total number of compounds in the MoA.

Image access and autocorrelation analysis was performed in Python, while results analysis and plotting were performed in R and Prism.

## Acknowledgements

We are grateful for the deposition of the annotated original images on Open Microscopy by the authors of the phenotypic screen (8) and for IDR API development.

## Author Contribution

Both authors contributed to each aspect of the manuscript.

## Competing interests

R.A.D. is an employee of Siren Pharmaceuticals LLC. R.A.D. and J.M.D. own equity in Siren Pharmaceuticals LLC.

